# Immune gene expression covaries with gut microbiome composition in stickleback

**DOI:** 10.1101/2020.08.04.236786

**Authors:** Lauren Fuess, Stijn den Haan, Fei Ling, Jesse N. Weber, Natalie C. Steinel, Daniel I. Bolnick

**Author notes:** Corresponding Author:* Lauren Fuess.

## Abstract

Commensal microbial communities have immense effects on their vertebrate hosts, contributing to a number of physiological functions as well as host fitness. In particular, host immunity is strongly linked to microbiota composition through poorly understood bi-directional links. Gene expression may be a potential mediator of these links between microbial communities and host function. However few studies have investigated connections between microbiota composition and expression of host immune genes in complex systems. Here we leverage a large study of laboratory-raised fish from the species *Gasterosteus aculeatus* (three-spined stickleback) to document correlations between gene expression and microbiome composition. First, we examined correlations between microbiome alpha diversity and gene expression. Our results demonstrate robust positive associations between microbial alpha diversity and expression of host immunity. Next, we examined correlations between host gene expression and abundance of microbial taxa. We identified 15 microbial families that were highly correlated to host gene expression. These families were all tightly correlated to host expression of immune genes and processes, falling into one of three categories: those positively correlated, negatively correlated, and neutrally related to immune processes. Furthermore, we highlight several important immune processes that are commonly associated with abundance of these taxons, including both macrophage and B cell functions. Further functional characterization of microbial taxa will help disentangle the mechanisms of the correlations described here. In sum, our study supports prevailing hypotheses of intimate links between host immunity and gut microbiome composition.

## INTRODUCTION

Diverse communities of commensal microbiota are associated with a range of vertebrate organ systems, such as the respiratory tract (Beck et al., 2012), skin (Grice and Segre, 2011), and digestive tracts (Backhed et al., 2005). Each of these distinct communities can contribute to host physiological function and fitness (e.g., (Jandhyala et al., 2015; Marchesi et al., 2016), but can cause pathology when disrupted (e.g., (Cho and Blaser, 2012). Conversely, host function, including the host immune system, also has effects on the composition of the microbiota, maintaining mutualists to avoid dysbiosis while also eliminating disease-causing pathogens (Belkaid and Hand, 2014; Levy et al., 2017). Because host immune and physiological functions often entail changes in gene expression, these reciprocal interactions between host function and microbiome structure may be revealed by examining host gene expression (Broderick et al., 2014; Richards et al., 2019). Preliminary studies suggest that changes in microbiome can affect host gene expression (Richards et al., 2019) and vice versa (Blekhman et al., 2015; Org et al., 2015). Still, much of what is known regarding cross-talk between host gene expression and microbiota composition has been learned using reduced, or single taxa microbial models in a handful of host species (Camp et al., 2014; Geva-Zatorsky et al., 2017; Sommer et al., 2015). A potentially valuable next step would be to examine transcriptome-wide associations with variation in the entire gut microbial community. However, such studies are lacking because until recently transcriptomic analyses were too expensive to allow sufficient statistical power (Lohman et al 2017).

Preliminary evidence suggests that host immunity in particular is closely linked to microbiota composition, likely through complex feedback networks (Belkaid and Hand, 2014; Zheng et al., 2020). Vertebrates’ mutualist bacteria promote key immune tolerance and regulatory pathways in a variety of organ systems (Belkaid and Hand, 2014). These include innate immunity in skin (Lai et al., 2010; Wanke et al., 2011), response to influenza in the respiratory tract (Ichinohe et al., 2011), and development of gut associated lymphoid tissue (GALT) and regulatory T cells in the intestines (Jandhyala et al., 2015). Removal of or changes in these microbes compromises host immunity (Reikvam et al., 2011) or can lead to auto-immune disorders (Jangi et al., 2016; Scher and Abramson, 2011). Furthermore, studies have shown the importance of commensal bacteria in regulating and educating adaptive immunity (Honda and Littman, 2016; Zhao and Elson, 2018), as well as contributing to development and homeostasis of innate immune cells (Balmer et al., 2014; Deshmukh et al., 2014; Khosravi et al., 2014). Conversely, on the part of the host, numerous immunological systems function in regulating microbiota composition, including physical barriers (i.e. host-secreted mucus layers; (Okumura and Takeda, 2018; Schroeder, 2019), host-produced anti-microbial compounds (Salzman, 2010), recognition molecules (i.e. PRRS; (Chu and Mazmanian, 2013), and associated signaling; (Kubinak et al., 2015) and effector responses (i.e. secretion of antibodies; (Nakajima et al., 2018). Despite considerable preliminary knowledge regarding cross-talk between host immunity and microbiota composition, the mechanisms of this feedback, particularly in regards to the roles of host gene expression, are not well described.

Advances in transcriptomics (RNAseq), have allowed for improved understanding of host function, including immunity and immune response, in diverse systems (Dheilly et al., 2014). These advancements can allow for the expansion of work investigating bidirectional interactions between microbiota and host immunity (hereafter, ‘microbe-immune feedbacks’) beyond existing laboratory model systems with simplified microbial compositions (Chevrette et al., 2019; Douglas, 2018; Macpherson and McCoy, 2015; Rosshart et al., 2017; Smith et al., 2007; Winans et al., 2017). Despite these technical advances, RNAseq has yet to be broadly applied to investigating microbe-immune feedbacks, particularly in complex contexts. A handful of studies have indicated correlation between microbiome composition and expression of immune genes (Donovan et al., 2014; Inoue et al., 2016). One such study screened a diversity of microbial species for their effects on host gene expression (whole transcriptome using Affymetrix arrays), demonstrating complex immunomodulatory effects of symbiotic microbes (Geva-Zatorsky et al., 2017). However these studies mostly document the effects of simplified microbiota or even monocultures. Only one study has examined more complex interactions, demonstrating strong associations between gene expression and microbiome composition in colonic epithelial cells, though these associations were limited to the localized colonic environment (Richards et al., 2019). Indeed, most studies examining feedback between microbiome composition and host gene expression have focused on localized gene expression, particularly in the gut epithelial tissue (Larsson et al., 2012; Qin et al., 2018). Consequently, we know little about the system-wide effects of microbiota composition on expression in distant immune-relevant tissues, and vice versa. Finally, few of these studies have examined the effects of genetic and environmental variation among hosts on these relationships.

Here, we report evidence for covariation between host gene expression and their gut microbiota, from a large sample of laboratory-bred threespine stickleback (*Gasterosteus aculeatus*), a small fish native to north temperate coastal marine and freshwater habitats. Like many vertebrates, individual stickleback harbor hundreds of microbial taxa (OTUs) in their intestines (Bolnick et al., 2014a; Bolnick et al., 2014b; Bolnick et al., 2014c; Ling et al., 2020). The composition of this microbiota differs dramatically between co-occurring individuals within a given natural population, and between populations (between neighboring lakes, adjacent lake and streams, or marine versus freshwater; (Bolnick et al., 2014a; Bolnick et al., 2014b; Bolnick et al., 2014c; Milligan-Myhre et al., 2016; Rennison et al., 2019; Smith et al., 2015). For example, in one survey of a single natural population of stickleback, proteobacteria ranged from less than 5% to over 95% of the microbial community depending on the individual host. This dramatic among-individual variation is associated with variation in diet, sex, genotype (at MHC and other autosomal loci), helminth infection status, and interactions between these factors (Bolnick et al., 2014a; Bolnick et al., 2014b; Bolnick et al., 2014c; Ling et al., 2020). Similar among-individual variation is observed within laboratory stocks of stickleback, whose microbiota is partly but not fully overlapping with the taxa seen in wild populations (Bolnick et al., 2014c).

Here, we seek to test whether this among-individual variation in gut microbiota composition in the laboratory is associated with individuals’ immune gene expression. Using transcriptomic data generated from head-kidneys (primary immune organ), we document correlations between immune gene expression and both microbiome diversity, and proportion of key microbial families. Our results are some of the first to describe links between broad host immune functioning and microbiome structure in a non-mammalian vertebrate.

## METHODS

### Experimental Design

Full details of data collection can be found in Ling et al. (Ling et al., 2020) and Fuess et al. (2020). Briefly, we collected reproductively mature fish from two lakes on Vancouver Island, British Columbia (Roberts and Gosling Lakes). Eggs were removed from gravid females and fertilized using sperm from testes collected from males from the same lake (pure F1s), or from the other lake (F1 hybrids). Eggs were shipped back to the University of Texas at Austin, hatched, and reared to reproductive maturity. The resulting adults were again artificially crossed to generate F2 hybrids consisting of intercrosses (F1×F1 hybrids) or backcrosses (ROBxF1 or GOSxF1). The resulting generation was reared for 1 year, and then experimentally exposed to *S. solidus* cestodes, following standard procedures (Weber et al., 2017a; Weber et al., 2017b). Forty-two days post-exposure, fish were euthanized and data were collected for a number of phenotypic metrics including sex, mass, and infection status/load. We also dissected out head kidneys for immune transcriptomic analysis. In fish the head-kidney, or pronephros, is a primary immune organ functioning primarily as a lympho-myeloid compartment (Kum and Sekkin, 2011). As is indicated by the name, this structure is located in the cranial region of the fish, near the gills, separating it considerably from the gut. Guts were dissected using sterile protocols for microbiome composition analysis (Ling et al., 2020).

### Transcriptomic Analysis

RNA was extracted from one head kidney and sequencing libraries were generated following methods described in Fuess et al (Fuess et al., 2020). We extracted RNA from this organ using the Ambion MagMAX-96 Total RNA Isolation Kit, following a modified version of the manufacturer’s protocol. A DNA removal step was preformed using TURBO DNAse. RNA yield was quantified using a Tecan NanoQuant Plate. TagSeq RNA sequencing libraries were constructed using a modified version of methods described in Lohman et al. (Lohman et al., 2016), detailed in Fuess et al. (Fuess et al., 2020). Libraries were sequenced on a HiSeq 2500 at the Genomics Sequencing and Analysis Facility of the University of Texas at Austin.

Resulting sequencing reads were processed using the iRNAseq pipeline (Dixon et al., 2015). Reads were aligned to version 95 of the stickleback genome on Ensembl using Bowtie 2 software (Langmead and Salzberg, 2012), and any samples with less than 500,000 aligned reads were discarded (final N = 393). A matrix of normalized read counts was generated using the R package DESeq2 (Love et al., 2014). This normalized read count matrix was used for all subsequent analyses (correlations, path analyses, and WGCNA). Information about the resulting read counts per individual, and other metrics of transcriptome information are reported in Fuess et al. (2020).

### Gut Microbiota Analyses

Full details regarding sampling and analysis of gut microbiota composition can be found in Ling et al. (Ling et al., 2020). To summarize: DNA was extracted from the entirety of collected stickleback intestines (*n* = 693 fish) using MoBio Powersoil DNA Isolation Kits. From this data, 16S rRNA amplicons were generated for the V4 hypervariable regions. Sequencing was performed on an Illumina Miseq platform at the Genomic Sequencing and Analysis Facility at the University of Texas at Austin. Resulting reads were processed using standard procedures in the mothur software package (v.1.39.1; (Schloss et al., 2009). OTUs were identified using the UCLUST algorithm based on 97% similarity. Relative proportion of microbial taxa (calculated at the level of Family) was calculated as the proportion of total OTU reads from a sample representing a given family compared to the total number of OTUs for a sample. Data was rarefied to 2000 sequences and Chao1 alpha diversity was calculated using the R package phyloseq (McMurdie and Holmes, 2013). Information about the resulting number of microbial OTUs, counts, and read depth per individual are reported in greater detail in Ling et al. (2020), who examined the microbiota’s response to cestode infection and host genotype.

### Correlative Analyses

We tested for correlations between microbiome composition and host gene expression, All statistics were conducted in R (v. 3.6.1). First, we correlated gene expression of all expressed genes to alpha diversity using a Kendall’s rank correlation. Genes with p-values less than 0.05 were considered significant for further analyses. Next, to identify families of microbe that are highly associated with gene expression, we correlated gene expression of all expressed genes to the relative proportion of each microbial family, again using a Kendall’s rank correlation. This resulted in thousands of significant associations between families and genes, many of which may be false positives due to the exceptionally large number of tests run (due to considering the combinations of many transcripts, against many microbes). We concluded the most conservative approach would be to select the top 5% (approximate; ties accounted for) most significantly correlated families (15) and genes (1290), and consider only relationships between these two groups for further analyses.

To assess the effects of co-varying factors (i.e. successful cestode infection, sex, host genotype [cross]) and ensure that correlations were not the result of spurious covariate effects, we also conducted a path analysis using the R package *sem* (Fox, 2006). Potential co-variates which may have confounded relationships detected by the correlative analyses were included in the model: sex, infection, mass (log-transformed), and cross-direction. The full model structure can be found in the supplementary materials (**Supplementary Figure 1**).

### Gene Ontology Analyses

To determine the biological processes most correlated to microbiome diversity and composition, we conducted gene ontology analyses. We assessed enrichment of biological process GO terms, using the R script GO-MWU (Wright et al., 2015). To identify biological processes enriched as a result of variation in microbiome diversity, we conducted gene ontology enrichment analyses using the tau values for all significantly correlated transcripts; all other genes were assigned a value of 0. We used a similar approach for assessing biological process terms enriched in relation to relative family proportion for each of our families of interest. Gene ontology analyses were conducted independently for each family. Input for this analysis was a matrix comprised of tau values for all significant correlations between a given family and the top 5% of genes (all other genes assigned a value of 0).

### Co-expression Analyses

We used co-expression analyses to identify groups of co-expressed host genes that were significantly correlated with microbiome diversity or relative proportion of microbial families (using only the 15 most significantly correlated families identified previously). Co-expression analyses were run using the R package WGCNA (Langfelder and Horvath, 2008). We constructed a signed network using bicor analyses and the following parameters: soft power = 12; minimum module size = 30; deepSplit = 2; dissimilarity threshold = 0.2. The resulting network was correlated to microbiome diversity and relative family proportion using a bicor correlation. Modules with significant correlations (*p* < 0.05) were analyzed for enrichment of biological processes using gene ontology enrichment (GO-MWU; default parameters for analysis of WGCNA modules).

## RESULTS

### Correlations between gene expression and microbiome diversity

Correlative analyses revealed strong associations between microbiota diversity and host gene expression, including expression of putative immune genes. Alpha diversity of the gut-associated microbiota was significantly correlated to 1929 transcripts involved in a range of functions (**Supplementary File 1;** ~7.5% of all transcripts). These correlations were robust to experimental co-variates: path analysis revealed 1014 (52.5%) of these correlations remained significant when accounting for sex, infection, mass, etc. We will henceforth discuss all 1931 of the identified correlated transcripts. Of this total, 834 transcripts were positively correlated to microbial diversity and 1095 were negatively correlated to diversity. Many of the correlated transcripts were involved in different arms of immunity (**Table 1**). Genes significantly correlated to microbiome diversity were significantly enriched for 11 biological process GO terms, 10 positively (i.e. over-represented processes which are increasing as a result of increased diversity of the microbiota based on tau values) and 1 negatively (i.e. over-represented processes whose expression is lower in fish with high microbial diversity; **Figure 1**). This included two terms involved in immunity that were positively correlated to diversity: *positive regulation of interleukin-12 production* and *common myeloid progenitor cell proliferation*. Genes that significantly contributed to enrichment of these two terms included Receptor-type tyrosine-protein kinase FLT3 (ENSGACT00000004059), Toll-like receptor 9 (ENSGACT00000013443), Tumor necrosis factor receptor superfamily member 5 (ENSGACT00000014780), Interferon regulatory factor 8 (ENSGACT00000021099), and Peregrin (ENSGACT00000001616). In contrast, the immune-related term *regulation of macrophage inflammatory protein 1 alpha production*, and associated genes, were expressed at lower levels in fish with more diverse gut microbiota. Significant genes included in this term were: Pyrin (ENSGACT00000027215), High mobility group protein B1 (ENSGACT00000027215), and Transient receptor potential cation channel subfamily V member 4 (ENSGACT00000012089). In sum, microbial diversity was positively correlated to the development of immune cells and regulation of IL-12, but negatively correlated to inflammatory processes.

**Table 1:** Examples of immune genes that were significantly correlated to diversity of gut-associated microbiota. A full list of significantly correlated genes can be found in **Supplementary File 2**.

| Gene | Ensembl ID | Immune Function | tau value | p-value |
| --- | --- | --- | --- | --- |
| C-C motif chemokine 4 | ENSGACT00000000554 | Chemoattractant (NK cells, monocytes, etc) | 0.197 | 4.76E-07 |
| Interferon regulatory factor 4 | ENSGACT00000021099 | Transcriptional activator (antiviral) | 0.152 | 9.96E-05 |
| Complement C3 | ENSGACT00000026259 | Complement cascade (innate immunity) | 0.133 | 7.65E-04 |
| Eosinophil peroxidase | ENSGACT00000022724 | Anti-bacterial activity | -0.175 | 7.68E-06 |
| Interleukin-11 | ENSGACT00000013923 | Hematopoietic stem cell proliferation | -0.152 | 0.00134 |
| NLR family CARD domain-containing protein 3 | ENSGACT00000001559 | Negatively regulates innate immunity | -0.143 | 0.00272 |

**Figure 1:**
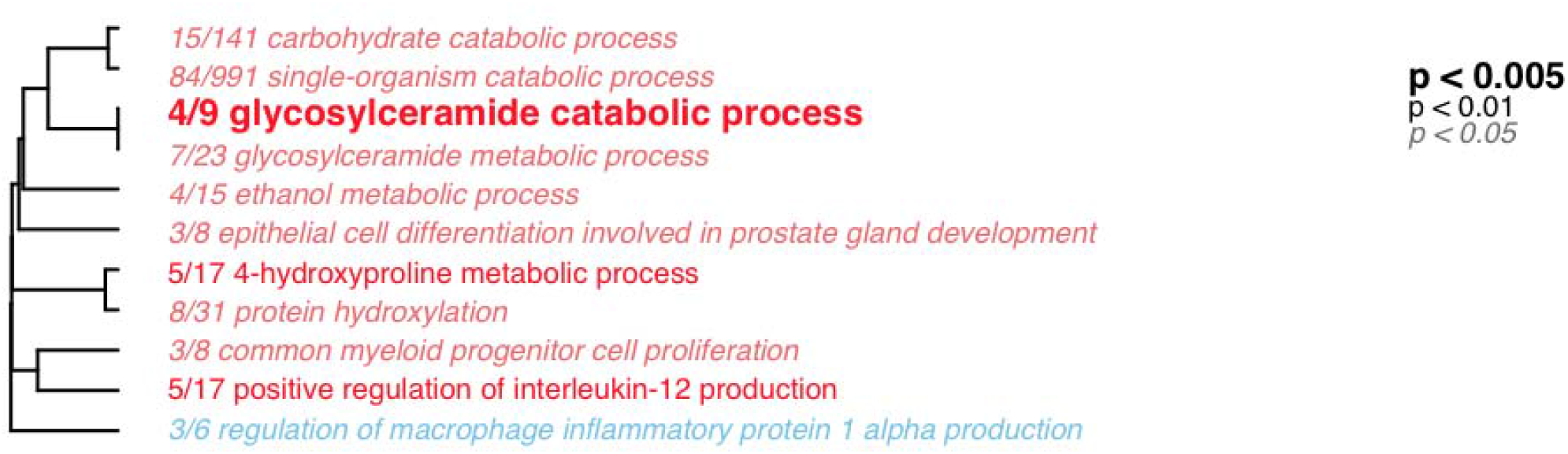
Hierarchical clustering of significantly enriched biological process gene ontology terms associated with genes significantly correlated to microbial diversity. Terms in red are positively enriched, terms in blue are negatively enriched. Font style indicates level of significance.

Coexpression analyses revealed strong associations between microbiota diversity and host gene expression, and immunity in particular. The resulting network comprised of 10 modules, plus a “grey” module containing transcripts that did not fit into any existing modules. These modules ranged in size from 44 to 18,227 transcripts. The largest of these modules (turquoise) likely represent groups of housekeeping genes with lowly variable expression. Two modules were significantly positively correlated to microbial diversity: yellow (r = 0.11, p=0.03) and magenta (r = 0.12, p=0.02; **Supplementary Figure 2**). The yellow module (1344 transcripts) was significantly enriched for 76 biological process gene ontology terms involved in a wide diversity of processes, indicating its roles in basic cellular homeostasis. Some of the largest groups of these terms included those involved in translational initiation, cytosolic transport, cellular component biogenesis, and electron transport chain (**Supplementary Figure 3**). The biological meaning of this module is ambiguous. In contrast, the much smaller magenta module (48 transcripts) was enriched for 36 biological process gene ontology terms, of which 20 were related to immunity and defense (**Figure 2**). Thus we concluded that the magenta module consists of co-regulated genes predominately involved in immune function. Enriched terms included those involved in interferon production (*positive regulation of type-I interferon production, positive regulation of interferon-alpha production, interferon-gamma-mediated signaling pathway* etc), and cytokine signaling (*cytokine-mediated signaling pathway, regulation of cytokine production,* etc), as well as other general immune GO terms (*immune response, immune effector process, innate immune response*, etc.). Thus, fish with more diverse microbiota generally exhibited higher co-expression of these categories of immune genes. Consequently, coexpression analyses indicate strong positive association between host gene expression, and a diverse network of genes involved in immunity, with emphasis on interferon and cytokine signaling.

**Figure 2:**
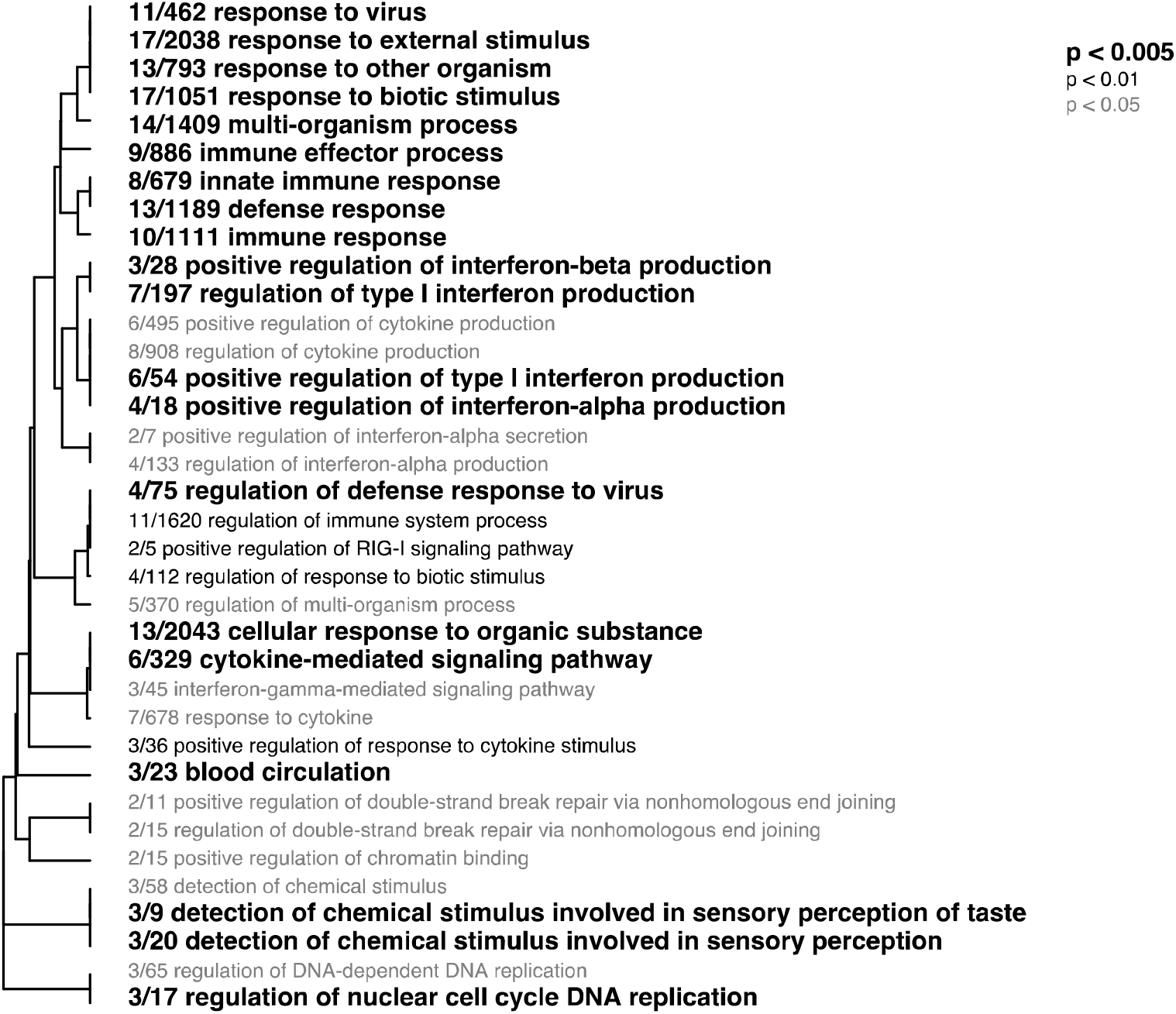
Hierarchical clustering of significantly enriched biological process gene ontology terms associated with genes in the magenta module (positively associated with microbial diversity). Font style indicates level of significance.

### Correlations between gene expression and relative abundance of specific taxa

Initial analysis of correlations between microbial families and host gene expression identified 507,317 significant associations out of 7,893,582 possible pairwise correlations between the relative abundance of a given family and a specific gene (~6.4% of total correlations run, slightly but very significantly more than the 5% expected from Type II error alone). We took a conservative approach and further examined only correlations between approximately the top 5% (approximate) most correlated microbial families (15) and genes (1297). Combined, these families were correlated to a total of 1263 of the 1297 top correlated genes (**Figure 3; Supplementary File 2**). Again most of these filtered relationships were robust to covariates (**Table 2**), thus herein we will discuss all significant results from the initial analysis.

**Table 2:**
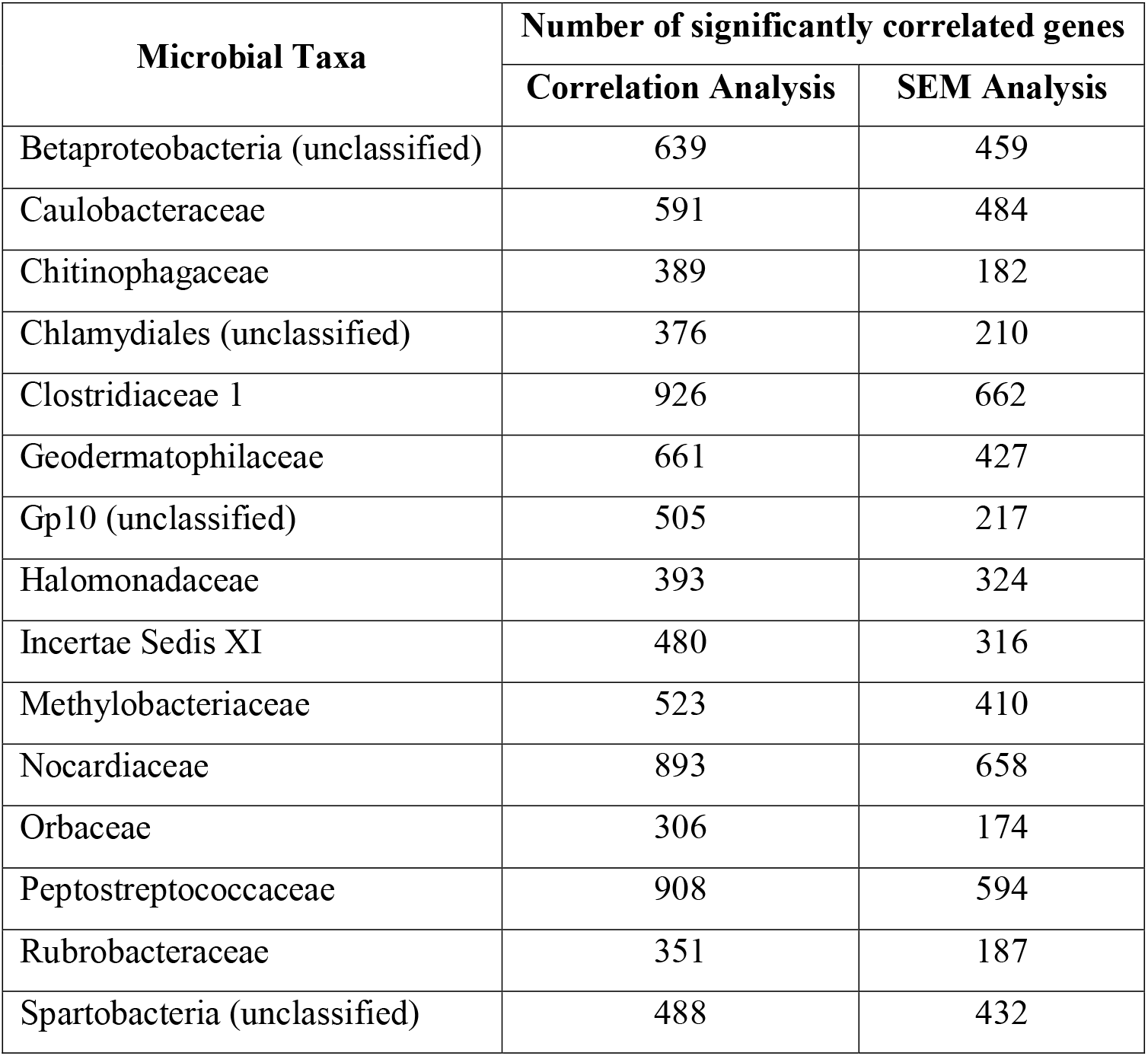
Summary of statistically significant correlations between the top 5% most correlated families and genes. Also listed are the number of correlations which remain significant following covariate analyses using structural equation modeling (SEM analysis).

**Figure 3:**
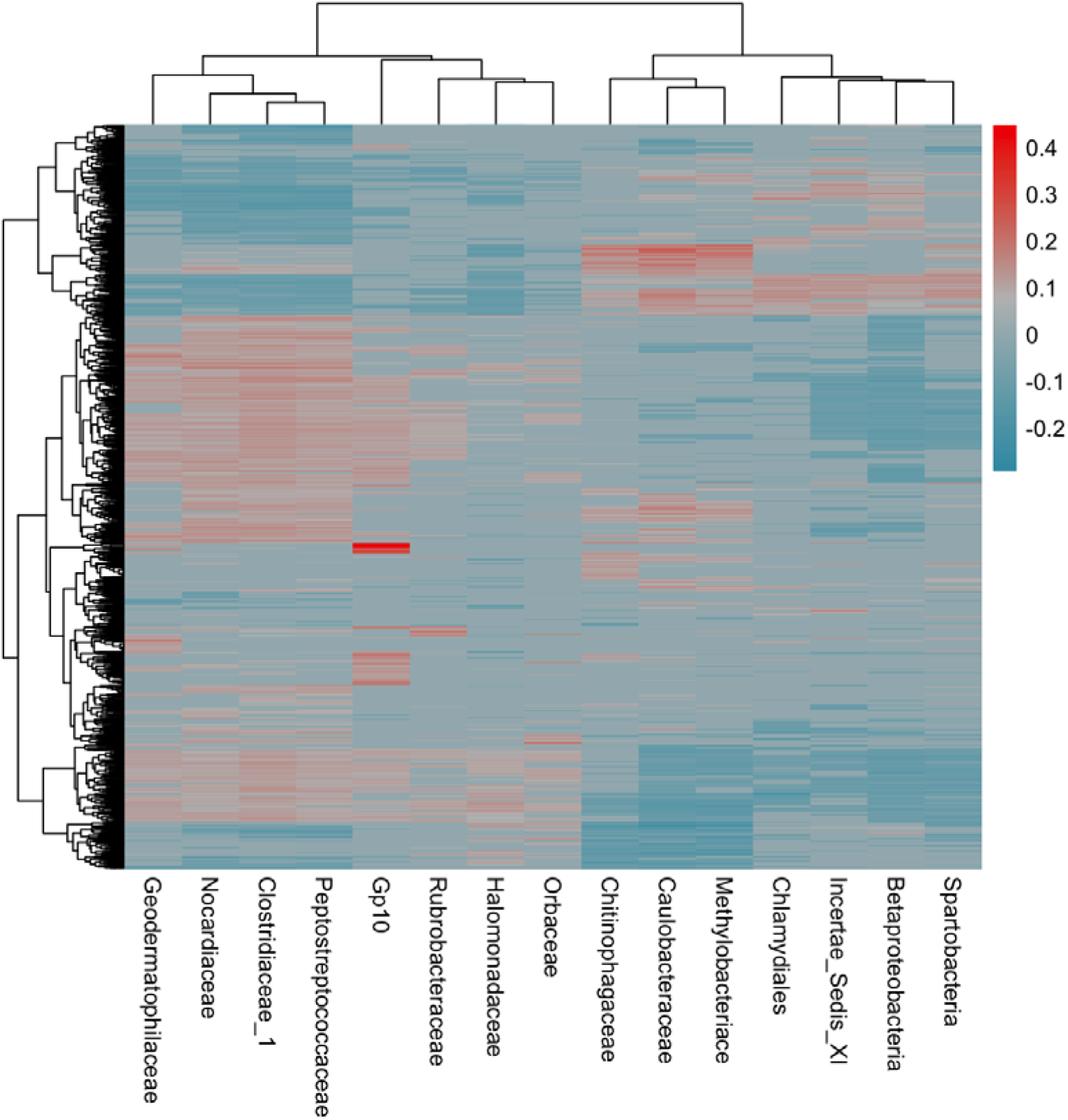
Heatmap of correlations between families and genes of interest. Values shown are Kendall’s tau for each correlation. Grey fill indicates non-significant correlations. Rows and columns are hierarchically clustered using default parameters.

Gene ontology enrichment of genes correlated to each family revealed significant patterns of enrichment of immune processes. The 15 families fell into one of three categories based on gene ontology enrichment analyses: positively immune associated, neutrally immune associated, or negatively immune associated (**Figure 3; Supplementary File 3**). Families such as Rubrobacteraceae, Orbaceae, and Halomonadaceae showed significant negative associations with immunity. Genes correlated to Rubrobacteraceae and Halomonadaceae abundance were significantly negatively enriched for immune-associated biological process GO terms such as *lymphocyte aggregation, response to interleukin-15,* and *myeloid lymphocyte migration*. Similarly, Orbaceae was significantly correlated to genes negatively enriched for immune terms including: *pro-B cell differentiation, positive regulation of interferon-alpha secretion,* and *hummoral immune response*. In contrast, abundance of six microbial taxa, including Caulobacteraceae and Chlamydiales were positively associated with immune processes. Genes correlated to these taxa were positively enriched for immune-associated biological process GO terms such as *positive regulation of B cell differentiation*, *positive regulation of macrophage activation, positive regulation of interleukin-12 production,* and *myeloid progenitor cell differentiation*. Six microbial families had mixed (i.e. neutral) associations with biological processes (**Figure 4**). Based on enrichment analysis of the correlative results, abundance of microbial taxa can have complex effects on host gene expression, dependent on the taxon identity.

**Figure 4:**
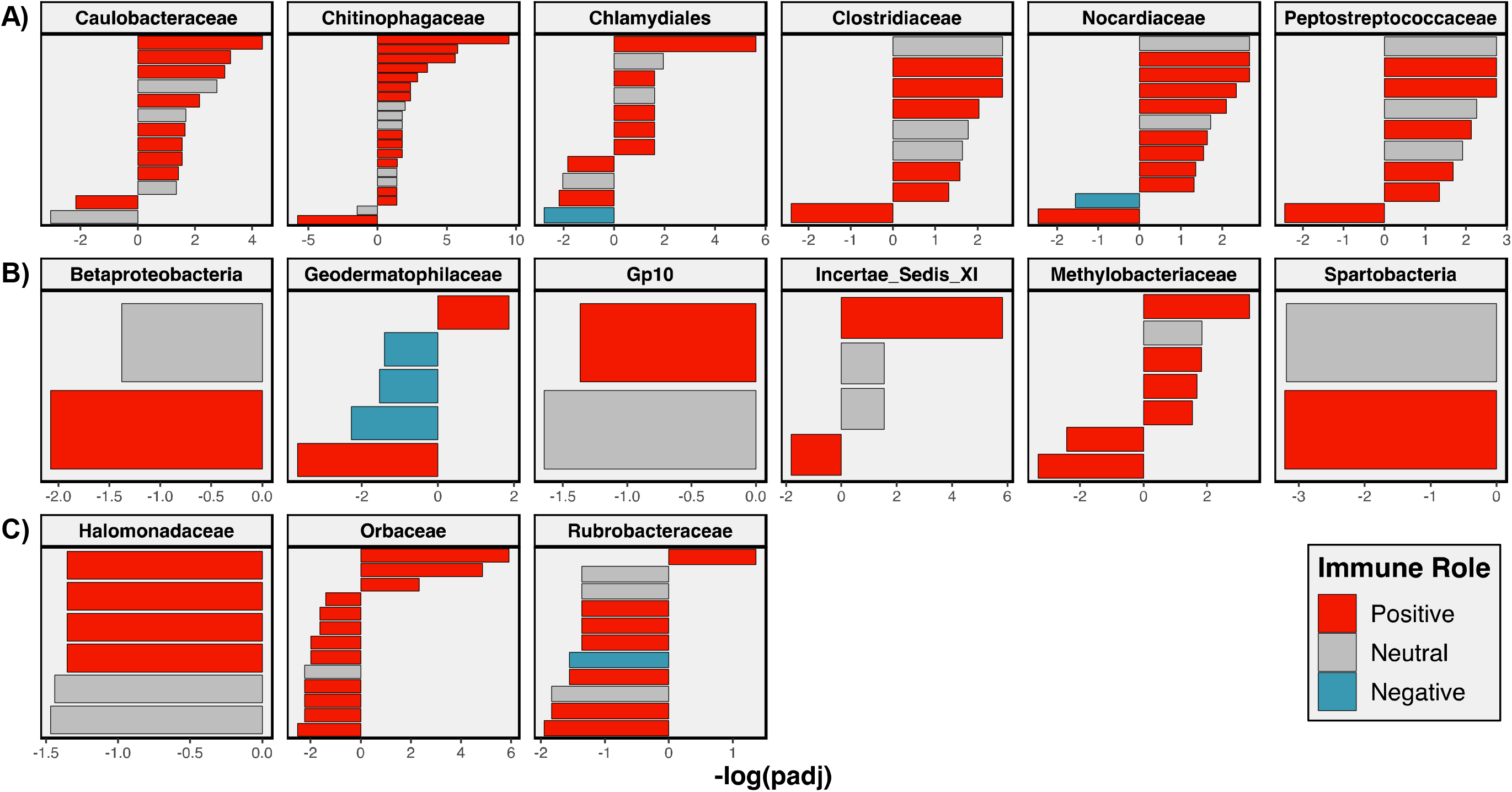
Significance and enrichment of immune associated biological process terms associated with each of the 15 microbial families of interest. Families are classified into three groups: **A)** families negatively associated with immunity, **B)** families neutrally associated with immunity, and **C)** families positively associated with immunities. Each bar indicates a significantly enriched biological GO term; color indicates the association of the term with immunity (red are terms which positively contribute to immune functions, blue are negative terms, and grey are neutral); direction of bar indicates positive or negative enrichment; magnitude of bars indicates the negative log of adjusted p-value.

Nine of ten WCGNA co-expression modules were correlated to one or more of the 15 identified microbial families of interest (**Supplementary Figure 2**). The magenta module, the immune functioning module significantly correlated to diversity, was only correlated to abundance of Spartobacteria. Of the nine modules that were correlated to one or more family of interest, most had broad biological functions. Exceptions to this included the pink module, which was enriched primarily for biological processes involved in tetrapyrrole metabolism and inorganic anion transport. The pink module was significantly positively correlated to Betaproteobacteria (*r* = 0.17, *p* = 0.0009) and significantly negatively correlated to Chitinophagaceae (*r* = 0.11*, p* = 0.03) and Peptostrepococcaceae (*r* = −0.12, *p* = 0.01). Additionally, the red module, which was enriched for numerous terms involved in wound healing, defense, and immune response, was positively correlated to both Chitinophagaceae (*r* = 0.11, *p* = 0.04) and Orbaceae (*r* = 0.11, *p* = 0.03) Co-expression analyses suggest that microbial families are related to broad networks of host gene expression in complex, family-dependent patterns.

## DISCUSSION

### Microbiome diversity is positively associated with host immunity

Numerous studies, mostly of laboratory mice, indicate that vertebrates’ microbiome composition is both a function of their immune genotype and phenotype (Blekhman et al., 2015; Bonder et al., 2016; Maynard et al., 2012; Oh et al., 2013), and in turn can modify host immune development and response (Chang et al., 2014; Maynard et al., 2012; Zhang et al., 2019). Despite this, the mechanisms of these relationships, as well as patterns of microbe-immune feedbacks in more complex and diverse systems are not known. Here we provide some of the first evidence of the roles of host gene expression in mediating microbe-immune feedbacks in a non-mammalian system. Based on existing evidence, we expect correlations between gut microbiota composition and host immunity, as measured by gene expression in immunological tissues. This should be true even when examining tissues that are anatomically separated from the organ(s) containing the microbiota, because localized interactions in the gut may alter system-wide immune traits. Consistent with this expectation, we find that alpha diversity of *G. aculeatus* gut micobiota was significantly correlated to expression of a large number of host genes expressed in the head kidney, many of which had functions in immunity. Co-expression analysis further confirmed strong associations between a broad diversity of immune componenss, and microbial diversity. Our results collectively confirm that, broadly speaking, host immunity (measured by transcriptomics) and microbiome diversity are positively associated. Furthermore, our results highlight the far-reaching effects of gut microbiome on host immunity: gut microbiome composition was correlated with gene expression broadly, and expression of immune genes especially, in the anatomically distant head kidney (located cranially near the gills). Furthermore, correlations between gut microbiota and gene expression in head kidney tissue indicate significant cross-talk between immune cell development and microbiota composition, as the head kidney is primarily involved in the development of a range of immune cells (Kum and Sekkin, 2011). Consequently, our findings suggest complex effects of gut microbiome on the development of host immune cells, and consequently host immune function.

Gene ontology analysis of significantly correlated genes emphasized the relationship between microbiome diversity and specific aspects of immunity in *G. aculeatus*. *Positive regulation of interleukin-12 production* was positively associated with microbiome diversity. This is in accordance with past studies demonstrating effects of pro- and prebiotic compounds on interleukin-12 production (Kang and Im, 2015; Llewellyn and Foey, 2017; Shokryazdan et al., 2017). Furthermore, *common myeloid cell progenitor differentiation* was positively associated with microbial diversity. These associations are likely the result of effects of microbiome diversity on host functioning, rather than vice versa (i.e. host characteristics shaping microbiome diversity). Commensal bacteria increase the amount and division of myeloid progenitor cells (Thaiss et al., 2016). Furthermore increased myelopoiesis (Thaiss et al., 2016) and bone marrow myeloid cell abundance (Balmer et al., 2014) are positively associated with microbiome complexity.

Germ-free mice are also characterized by reduced myeloid-cell development (Thaiss et al., 2016). Our results indicate similar relationships between gut microbiota and development of immune cells in a major teleost hematopoetic organ, the head kidney. Finally, *regulation of macrophage inflammatory protein 1-α (MIP1-α)* was negatively associated with microbiome diversity. MIP1-α is an inflammatory chemokine (Kang et al., 2011). Inflammation is well known to be linked with dysbiosis of the microbiome (Levy et al., 2017); reduced microbial diversity is associated with many inflammatory conditions of the gut (Levy et al., 2017; Manichanh et al., 2006). In sum, our analyses revealed significant connections between microbiome diversity and host immunity, though the mechanism and direction of causation of these relationships requires further study.

### Microbial taxa have opposing effects on host immune gene expression

In addition to highlighting significant microbe-immune feedbacks associated with diversity of the gut microbiota, our study also provides substantial initial evidence of the effects of specific microbial taxa on host gene expression and immunity. Abundance of specific microbial taxa was correlated to a wide array of host genes, with functions in a diversity of biological processes. Indeed, co-expression analysis demonstrated strong associations between abundance of particular microbial families and modules involved in broad host functioning. The exact nature of these associations varied among microbial families, with certain groups of microbial families displaying opposing trends of correlations to both individual genes, and co-expression modules. Specific microbial taxa, are known to effect broad host functioning (Hooper et al., 2001; Jandhyala et al., 2015; Pedersen et al., 2016; Rowland et al., 2018). Our results are among the first to highlight the complex nature of specific host-microbe interactions, the importance of host gene expression in mediating these interactions, and the effects of these relationships on a diversity of functions. Here we will specifically focus on variation in association between microbial taxa and host immunity.

Gene ontology analysis of associations between families and genes of interest revealed clear patterns of microbial family-host immune associations. Microbial taxa fell into three groups: those positively correlated, negatively correlated, or neutrally associated with immunity. It is known that certain groups of commensal bacteria, such as segmented filamentous bacteria (Ivanov et al., 2009), are positively associated with aspects of host immunity, while others, including *Bacteroides fragilis* (Mazmanian et al., 2008), induce a protective tolerogenic responses, suppressing immunity (Geva-Zatorsky et al., 2017). Furthermore, some pathogenic bacteria, including *Bacillus anthracis* and *Shigella*, may suppress host immunity during infection (Hann and Rathjen, 2010; Reddick and Alto, 2014), while others will induce immune responses (Kumar et al., 2011; Kumar et al., 2013). However, often these relationships are highly context dependent, and the mechanisms of these relationships are poorly understood. Here we break down the observed relationships between specific microbial family abundance and host immunity. We specifically highlight the need for increased functional understanding of these taxa in order to improve mechanistic knowledge of the microbiome and its effect on host function.

Abundance of six different microbial taxa was significantly positively associated with expression of immune genes in our study. These families were: Caulobacteraceae, Chitinophagaceae, Chlamydiales, Clostridiaceae, Nocardiaceae, and Peptostreptococcaceae. Half of these families (Caulobacteraceae, Chlamydiales, and Nocardiaceae) are well described for their association with disease and other pathologies. Caulobacteraceae is a member of the phylum Proteobacteria, which is associated with numerous intestinal and extra-intestinal diseases (Rizzatti et al., 2017). Specifically, members of the family Caulobacteraceae have been associated with pneumonia in HIV patients (Cribbs et al., 2016). Members of another commonly pathogenic taxon, Chlamydiales, were also positively associated with immunity. Microbes from this family elicit a strong innate immune response to infection (Rusconi and Greub, 2011) and are capable of manipulating that response (Mehlitz and Rudel, 2013; Rusconi and Greub, 2011). Finally, members of the family Nocardiaceae were also positively associated with the expression of immune-associated genes. This family is composed primarily of environmental and soil microbes, but some members are opportunistic pathogens (Goodfellow, 2014). For example, nocardiosis in fish is induced by microbes from this family, and resulting in increased cytokine responses (Tanekhy et al., 2010). In sum, positive associations between these taxa and expression of host immune genes is most likely indicative of the pathogenic nature of these microbes, inducing a host immune response.

The remaining three families that were positively associated with host immunity, Chitinophagaceae, Clostridiaceae, and Peptostreptococcaceae, have diverse roles in both environmental and host-associated microbial communities. Members of the family Chitinophagaceae are often described as components of the commensal microbiota of aquatic species, including lampreys (Li et al., 2016), and aquatic amphibians (Walke et al., 2015). These microbes are capable of degrading chitin (Rosenberg, 2014), perhaps explaining their capability to inhibit growth of the common fungal amphibian pathogen, *Batrachochytrium dendrobatidis* (Walke et al., 2015). Beyond this beneficial association within amphibians, little more is known regarding function of Chitionphagaceae in vertebrate commensal microbial communities. The family Clostridiaceae is a diverse taxon that includes a group of bacteria known as segmented filamentous bacteria (SFB). These microbes have broad effects on host immunity, including enhancing Th17 cell responses (Ivanov et al., 2009), and promoting increased IgA production in mice (Klaasen et al., 1993). Some SFB may confer protection against pathogenic microbes (Ivanov et al., 2009). Finally, Peptostreptococcaceae is a poorly described, yet diverse microbial taxon, which is often associated with the vertebrate gut microbiome. Much of what is known regarding this family is based upon extensive research regarding a single representative species, *Clostridium difficile* (Navaneethan et al., 2010). However this family is immensely diverse (Slobodkin, 2014), necessitating further functional study to understand broader associations of members of this taxon with host immunity. Some preliminary studies have suggested negative associations between Peptostreptococcaceae abundance and risk factors for autoimmune disorders (Russell et al., 2019). Still the directionality of this relationship remains unclear. It is possible that positive association between host immune function and Peptostreptococcaceae abundance is due to aberrant removal of Peptostreptococcaceae in hosts suffering from immune dysfunction (Russell et al., 2019). Alternatively, Peptostreptococcaceae could have some unknown function in the development of host immunity, the absence of which would result in development of autoimmune disorders. Indeed, further functional classification and controlled mechanistic studies will prove fruitful in understanding positive associations between the taxa identified here and host immune functioning.

In contrast to those taxa identified as positively associated with host immune gene expression, three families could be classified as negatively associated with host immunity: Halomonadaceae, Orbaceae, and Rubrobacteraceae. All three of these microbial families are poorly described, and two (Halomonadaceae and Rubrobacteraceae) have been primarily described as environmental microbes. Members of the family Halomonadaceae are well described as halophiles (Oren, 2008); the roles of this family in microbiome composition are not well understood. Limited studies indicate changes in Halomonadaceae proportions are associated with certain pathologies (Huang et al., 2018; Vaziri et al., 2013), and there is some indication that some members may be pathogenic (Asea et al., 2002). Similarly, Rubrobacteraceae consists of thermophilic environmental microbes (Albuquerque and da Costa, 2014), though members have been isolated from the gut of termites (Lefebvre et al., 2009). In contrast, members of the family Orbaceae are known to be associated with animal commensal microbiota, though previously this taxon has only been described within the gut microbiota of bees (Billiet et al., 2017). Orbaceae is a common component of the bee microbiome and is negatively associated with colony productivity (Horton et al., 2015), and positively associated with parasite infection (Palmer-Young et al., 2019). Our study is the first, to our knowledge, to report presence of these microbes in the vertebrate gut microbiome. Further functional characterization of these three taxa is necessary to interpret their associations with host immunity in the *G. aculeatus* system.

Finally it is worth noting common trends in correlations between specific immune components and microbial family abundance. Several immune components were correlated to at least a third of the significant microbial families, as revealed by gene ontology analysis. Many of these components have also been linked to microbiome function or composition. Commonly correlated components included myeloid progenitor cell differentiation, regulation of interleukin-12 secretion/production, interferon-gamma production, pro-B cell differentiation, and positive regulation of macrophage activation. We have previously discussed the importance of both IL-12 and myeloid progenitor cells in the maintenance of gut-microbiome composition. The production of interferon-gamma (IFNγ) has been linked to microbiome processes. IFNγ has been both positively and negatively linked to various microbial components: IFNy production decreased in piglets treated with a probiotic bacteria (Hou et al., 2015), and immunomodulary compounds from other bacteria (Zvanych et al., 2014). In contrast, IFNγ^+^ CD8 T cells are induced by other commensal bacteria in human guts (Tanoue et al., 2019).

We observed frequent correlation between microbial family abundance and both B cell processes and macrophage activity. Microbiota are known to have profound effects on B cell processes including diversification, production of IgA, and differentiation of regulatory B cells (Rosser et al., 2014; Zhao and Elson, 2018). Furthermore, on the part of the host, B cell production of IgA in particular is essential to maintenance of gut microbiome composition by restricting commensal growth and maintaining a diverse composition (Palm et al., 2015). Similarly, in some teleost fish such as rainbow trout, IgT is known to play important roles in microbiome homeostasis (Xu et al., 2020). Similar bi-directional relationships are known to exist between macrophages and gut microbiota. Microbial metabolites such as butyrate can modulate and reduce macrophage activity to promote tolerance of commensal bacteria (Chang et al., 2014). Macrophages can also shape gut microbiota structure, potentially by discriminating between commensal and pathogenic microbes (Wang et al., 2019). In sum, the most striking patterns of correlation between immune components and microbial family abundance add to existing literature supporting the roles of these specific arms of immunity, and specific cell types, in microbiome maintenance.

## CONCLUSIONS

Here we document one of the first investigations of correlations between natural gut microbiome composition and host transcriptomic gene expression in a non-mammalian vertebrate. Our results detail extensive correlation between the host’s transcriptome and both diversity and proportion of specific microbial families. Notably, these associations exist despite spatial separation between the microbiota and the organ where we measure expression, highlighting the systemic changes induced by gut microbiota. Both diversity and microbial family proportion are strongly correlated with expression of a diversity of host immune genes. Associations between immunity and microbial diversity likely reflect both the effects of a healthy, diverse microbiome on host immune system, and the need for a robust immune system to maintain this diversity. Trends in correlation between abundance of specific microbial families and host immune gene expression identified groups both positively and negatively associated with host immunity. Many of these trends support previous studies from other systems. However, increased functional understanding of microbial taxa is needed to interpret these trends. In sum, our results highlight the immense interconnectivity between host gene expression and gut microbiome composition, specifically in regard to immune function. These results also highlight the utility of the transcriptomic tools in enabling studies of microbe-immune feedbacks in wild populations and non-model animals.

---

**Supplementary Figure 1:**
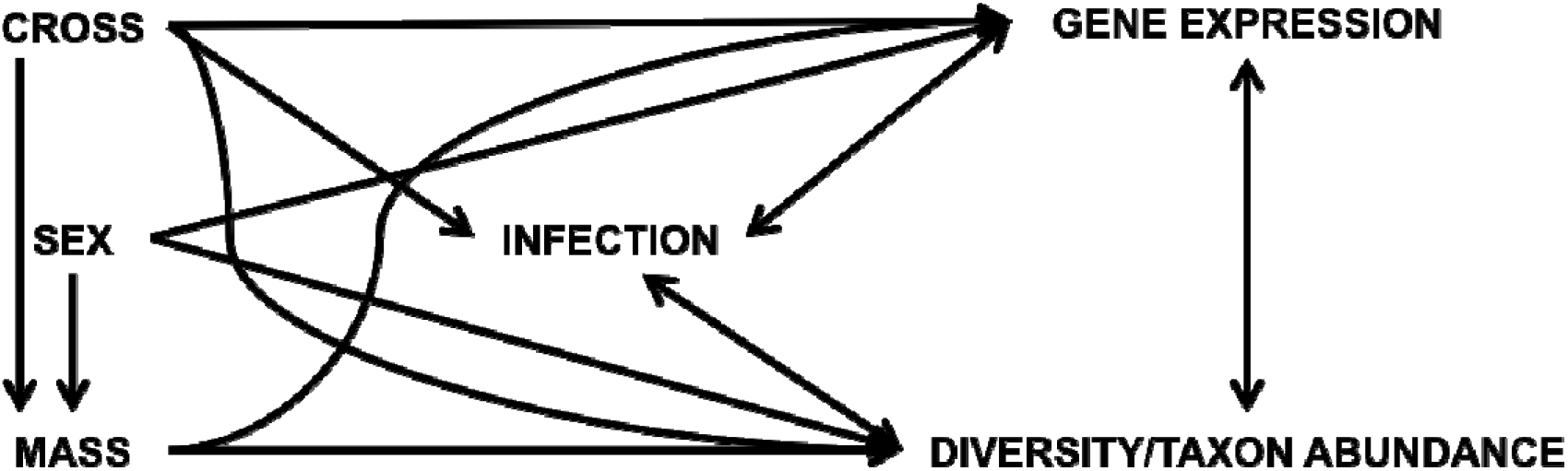
Schematic of structural equation model that was used to assess robustness of covariance between gene expression and microbial diversity or taxon abundance.

**Supplementary Figure 2:**
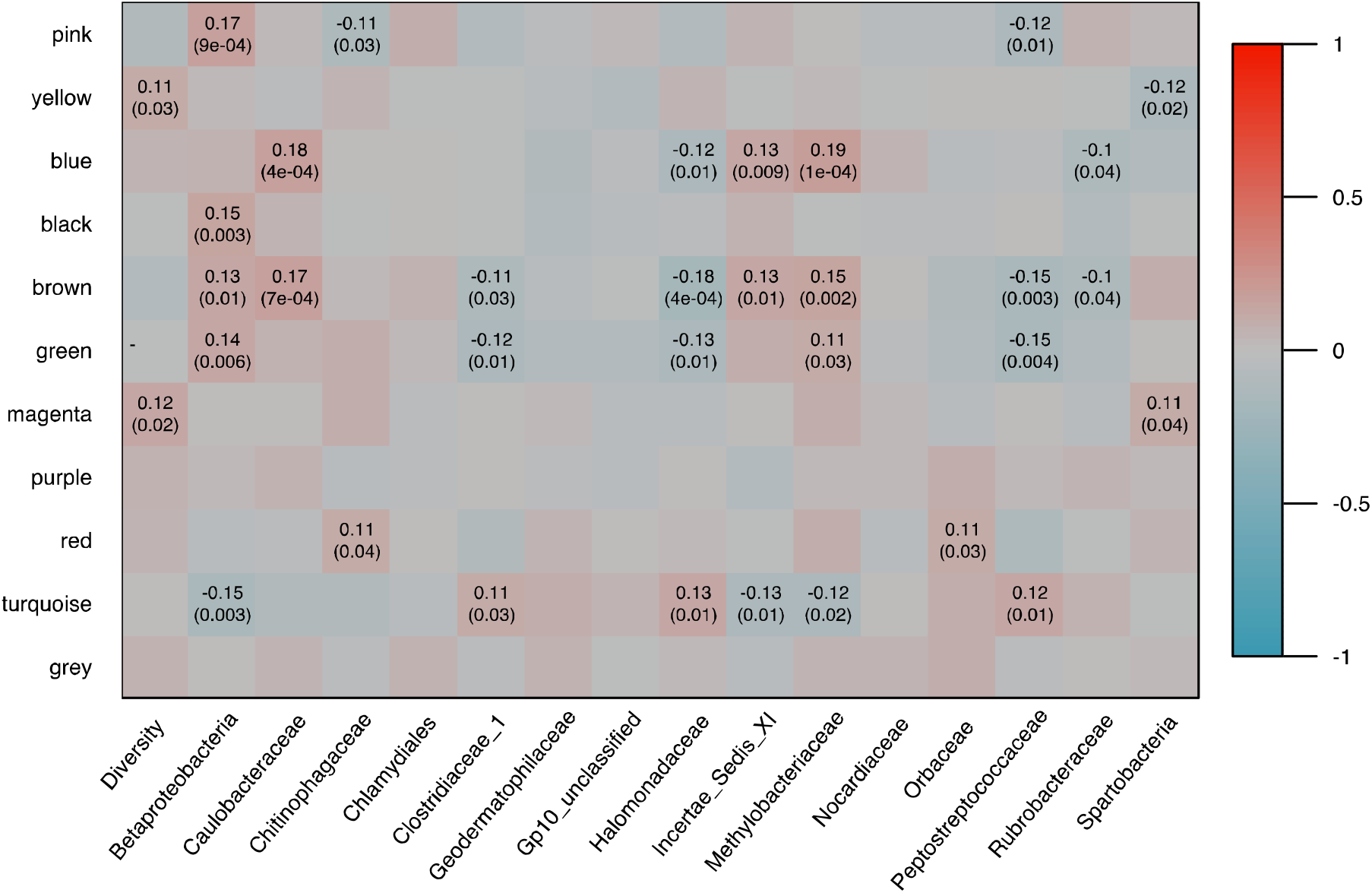
Tombstone plot displaying correlations between co-expression network modules and alpha diversity or microbial family diversity (for families of interest). Only significant values are displayed.

**Supplementary Figure 3:**
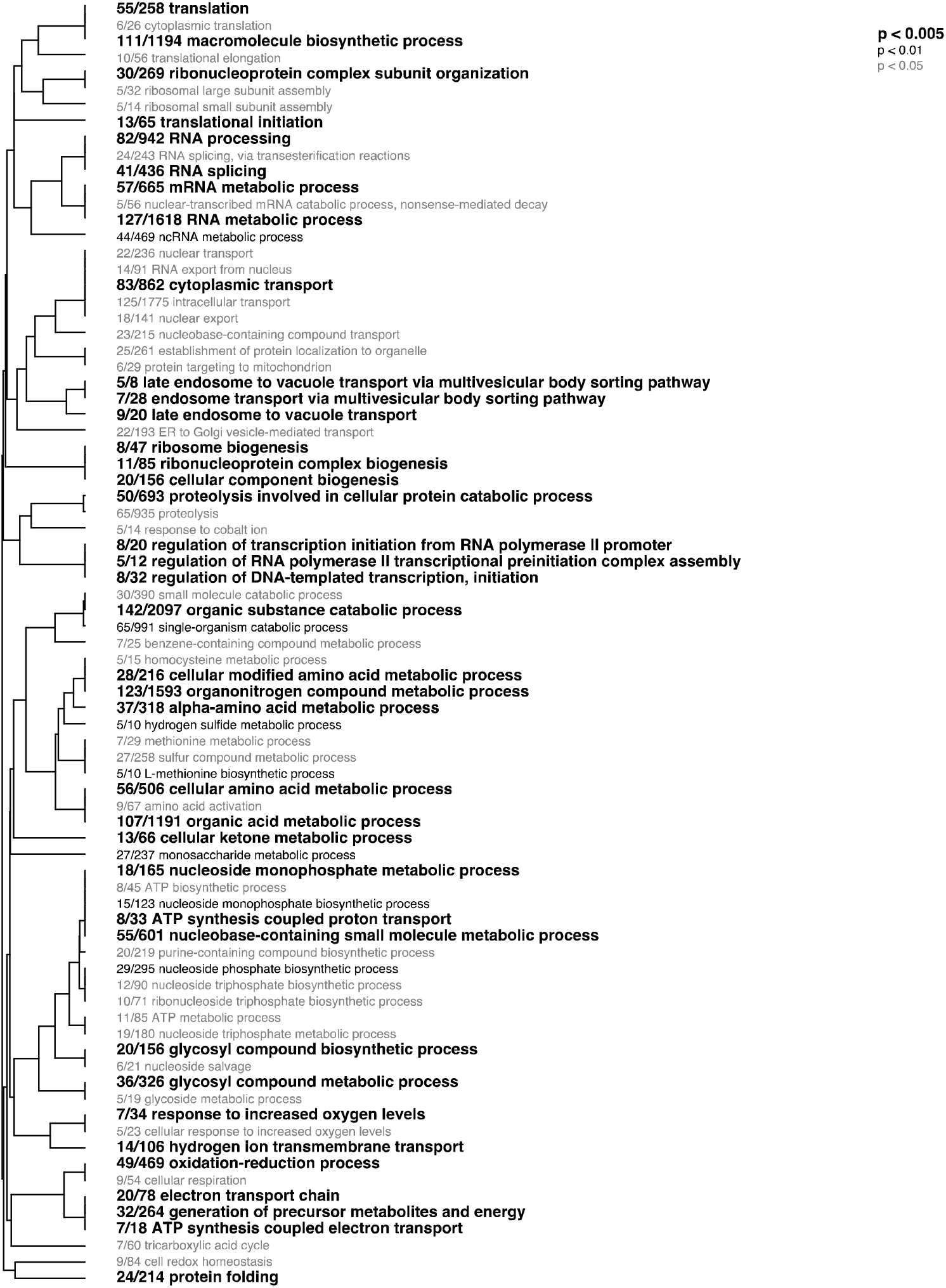
Hierarchical clustering of significantly enriched biological process gene ontology terms associated with genes in the yellow module (positively associated with microbial diversity). Font style indicates level of significance.

## References

Albuquerque, L., da Costa, M.S., 2014. The Family Rubrobacteraceae, in: Rosenberg, E., DeLong, E.F., Lory, S., Stackebrandt, E., Thompson, F. (Eds.), The Prokaryotes: Actinobacteria. Springer Berlin Heidelberg, Berlin, Heidelberg, pp. 861–866.

Asea, A., Rehli, M., Kabingu, E., Boch, J.A., Bare, O., Auron, P.E., Stevenson, M.A., Calderwood, S.K., 2002. Novel signal transduction pathway utilized by extracellular HSP70: role of toll-like receptor (TLR) 2 and TLR4. Journal of Biological Chemistry 277, 15028–15034.

Backhed, F., Ley, R.E., Sonnenburg, J.L., Peterson, D.A., Gordon, J.I., 2005. Host-bacterial mutualism in the human intestine. Science 307, 1915–1920.

Balmer, M.L., Schurch, C.M., Saito, Y., Geuking, M.B., Li, H., Cuenca, M., Kovtonyuk, L.V., McCoy, K.D., Hapfelmeier, S., Ochsenbein, A.F., Manz, M.G., Slack, E., Macpherson, A.J., 2014. Microbiota-derived compounds drive steady-state granulopoiesis via MyD88/TICAM signaling. Journal of immunology 193, 5273–5283.

Beck, J.M., Young, V.B., Huffnagle, G.B., 2012. The microbiome of the lung. Translational Research 160, 258–266.

Belkaid, Y., Hand, T.W., 2014. Role of the microbiota in immunity and inflammation. Cell 157, 121–141.

Billiet, A., Meeus, I., Van Nieuwerburgh, F., Deforce, D., Wackers, F., Smagghe, G., 2017. Colony contact contributes to the diversity of gut bacteria in bumblebees (Bombus terrestris). Insect Sci 24, 270–277.

Blekhman, R., Goodrich, J.K., Huang, K., Sun, Q., Bukowski, R., Bell, J.T., Spector, T.D., Keinan, A., Ley, R.E., Gevers, D., Clark, A.G., 2015. Host genetic variation impacts microbiome composition across human body sites. Genome biology 16, 191.

Bolnick, D.I., Snowberg, L.K., Caporaso, J.G., Lauber, C., Knight, R., Stutz, W.E., 2014a. Major Histocompatibility Complex class IIb polymorphism influences gut microbiota composition and diversity. Molecular ecology 23, 4831–4845.

Bolnick, D.I., Snowberg, L.K., Hirsch, P.E., Lauber, C.L., Knight, R., Caporaso, J.G., Svanbäck, R., 2014b. Individuals’ diet diversity influences gut microbial diversity in two freshwater fish (threespine stickleback and Eurasian perch). Ecology Letters 17, 979–987.

Bolnick, D.I., Snowberg, L.K., Hirsch, P.E., Lauber, C.L., Org, E., Parks, B., Lusis, A.J., Knight, R., Caporaso, J.G., Svanbäck, R., 2014c. Individual diet has sex-dependent effects on vertebrate gut microbiota. Nature communications 5, 4500.

Bonder, M.J., Kurilshikov, A., Tigchelaar, E.F., Mujagic, Z., Imhann, F., Vila, A.V., Deelen, P., Vatanen, T., Schirmer, M., Smeekens, S.P., Zhernakova, D.V., Jankipersadsing, S.A., Jaeger, M., Oosting, M., Cenit, M.C., Masclee, A.A., Swertz, M.A., Li, Y., Kumar, V., Joosten, L., Harmsen, H., Weersma, R.K., Franke, L., Hofker, M.H., Xavier, R.J., Jonkers, D., Netea, M.G., Wijmenga, C., Fu, J., Zhernakova, A., 2016. The effect of host genetics on the gut microbiome. Nat Genet 48, 1407–1412.

Broderick, N.A., Buchon, N., Lemaitre, B., 2014. Microbiota-induced changes in drosophila melanogaster host gene expression and gut morphology. mBio 5, e01117–01114.

Camp, J.G., Frank, C.L., Lickwar, C.R., Guturu, H., Rube, T., Wenger, A.M., Chen, J., Bejerano, G., Crawford, G.E., Rawls, J.F., 2014. Microbiota modulate transcription in the intestinal epithelium without remodeling the accessible chromatin landscape. Genome Res 24, 1504–1516.

Chang, P.V., Hao, L., Offermanns, S., Medzhitov, R., 2014. The microbial metabolite butyrate regulates intestinal macrophage function via histone deacetylase inhibition. Proc Natl Acad Sci U S A 111, 2247–2252.

Chevrette, M.G., Bratburd, J.R., Currie, C.R., Stubbendieck, R.M., 2019. Experimental Microbiomes: Models Not to Scale. mSystems 4, e00175–00119.

Cho, I., Blaser, M.J., 2012. The human microbiome: at the interface of health and disease. Nature Reviews Genetics 13, 260–270.

Chu, H., Mazmanian, S.K., 2013. Innate immune recognition of the microbiota promotes host-microbial symbiosis. Nature immunology 14, 668–675.

Cribbs, S.K., Uppal, K., Li, S., Jones, D.P., Huang, L., Tipton, L., Fitch, A., Greenblatt, R.M., Kingsley, L., Guidot, D.M., Ghedin, E., Morris, A., 2016. Correlation of the lung microbiota with metabolic profiles in bronchoalveolar lavage fluid in HIV infection. Microbiome 4, 3.

Deshmukh, H.S., Liu, Y., Menkiti, O.R., Mei, J., Dai, N., O’Leary, C.E., Oliver, P.M., Kolls, J.K., Weiser, J.N., Worthen, G.S., 2014. The microbiota regulates neutrophil homeostasis and host resistance to Escherichia coli K1 sepsis in neonatal mice. Nat Med 20, 524–530.

Dheilly, N.M., Adema, C., Raftos, D.A., Gourbal, B., Grunau, C., Du Pasquier, L., 2014. No more non-model species: the promise of next generation sequencing for comparative immunology. Developmental and comparative immunology 45, 56–66.

Dixon, G.B., Davies, S.W., Aglyamova, G.A., Meyer, E., Bay, L.K., Matz, M.V., 2015. Genomic determinants of coral heat tolerance across latitudes. Science 348, 1460–1462.

Donovan, S.M., Wang, M., Monaco, M.H., Martin, C.R., Davidson, L.A., Ivanov, I., Chapkin, R.S., 2014. Noninvasive molecular fingerprinting of host-microbiome interactions in neonates. FEBS Lett 588, 4112–4119.

Douglas, A.E., 2018. Which experimental systems should we use for human microbiome science? PLoS biology 16, e2005245–e2005245.

Fox, J., 2006. Structural Equation Modeling with SEM Package in R. Structural Equation Modeling-a Multidisciplinary Journal - STRUCT EQU MODELING 13, 465–486.

Fuess, L.E., Weber, J.N., den Haan, S., Steinel, N.C., Shim, K.C., Bolnick, D.I., 2020. A test of the Baldwin Effect: Differences in both constitutive expression and inducible responses to parasites underlie variation in host response to a parasite. bioRxiv, 2020.2007.2029.216531.

Geva-Zatorsky, N., Sefik, E., Kua, L., Pasman, L., Tan, T.G., Ortiz-Lopez, A., Yanortsang, T.B., Yang, L., Jupp, R., Mathis, D., Benoist, C., Kasper, D.L., 2017. Mining the Human Gut Microbiota for Immunomodulatory Organisms. Cell 168, 928–943 e911.

Goodfellow, M., 2014. The Family Nocardiaceae, in: Rosenberg, E., DeLong, E.F., Lory, S., Stackebrandt, E., Thompson, F. (Eds.), The Prokaryotes: Actinobacteria. Springer Berlin Heidelberg, Berlin, Heidelberg, pp. 595–650.

Grice, E.A., Segre, J.A., 2011. The skin microbiome. Nat Rev Microbiol 9, 244–253.

Hann, D.R., Rathjen, J.P., 2010. The long and winding road: virulence effector proteins of plant pathogenic bacteria. Cellular and molecular life sciences : CMLS 67, 3425–3434.

Honda, K., Littman, D.R., 2016. The microbiota in adaptive immune homeostasis and disease. Nature 535, 75–84.

Hooper, L.V., Wong, M.H., Thelin, A., Hansson, L., Falk, P.G., Gordon, J.I., 2001. Molecular analysis of commensal host-microbial relationships in the intestine. Science 291, 881–884.

Horton, M.A., Oliver, R., Newton, I.L., 2015. No apparent correlation between honey bee forager gut microbiota and honey production. PeerJ 3, e1329.

Hou, C., Liu, H., Zhang, J., Zhang, S., Yang, F., Zeng, X., Thacker, P.A., Zhang, G., Qiao, S., 2015. Intestinal microbiota succession and immunomodulatory consequences after introduction of Lactobacillus reuteri I5007 in neonatal piglets. PloS one 10, e0119505.

Huang, X., Ye, Z., Cao, Q., Su, G., Wang, Q., Deng, J., Zhou, C., Kijlstra, A., Yang, P., 2018. Gut Microbiota Composition and Fecal Metabolic Phenotype in Patients With Acute Anterior Uveitis. Invest Ophthalmol Vis Sci 59, 1523–1531.

Ichinohe, T., Pang, I.K., Kumamoto, Y., Peaper, D.R., Ho, J.H., Murray, T.S., Iwasaki, A., 2011. Microbiota regulates immune defense against respiratory tract influenza A virus infection. Proc Natl Acad Sci U S A 108, 5354–5359.

Inoue, R., Sakaue, Y., Sawai, C., Sawai, T., Ozeki, M., Romero-Pérez, G.A., Tsukahara, T., 2016. A preliminary investigation on the relationship between gut microbiota and gene expressions in peripheral mononuclear cells of infants with autism spectrum disorders. Bioscience, biotechnology, and biochemistry 80, 2450–2458.

Ivanov, II, Atarashi, K., Manel, N., Brodie, E.L., Shima, T., Karaoz, U., Wei, D., Goldfarb, K.C., Santee, C.A., Lynch, S.V., Tanoue, T., Imaoka, A., Itoh, K., Takeda, K., Umesaki, Y., Honda, K., Littman, D.R., 2009. Induction of intestinal Th17 cells by segmented filamentous bacteria. Cell 139, 485–498.

Jandhyala, S.M., Talukdar, R., Subramanyam, C., Vuyyuru, H., Sasikala, M., Nageshwar Reddy, D., 2015. Role of the normal gut microbiota. World journal of gastroenterology 21, 8787–8803.

Jangi, S., Gandhi, R., Cox, L.M., Li, N., von Glehn, F., Yan, R., Patel, B., Mazzola, M.A., Liu, S., Glanz, B.L., Cook, S., Tankou, S., Stuart, F., Melo, K., Nejad, P., Smith, K., Topçuolu, B.D., Holden, J., Kivisäkk, P., Chitnis, T., De Jager, P.L., Quintana, F.J., Gerber, G.K., Bry, L., Weiner, H.L., 2016. Alterations of the human gut microbiome in multiple sclerosis. Nature communications 7, 12015.

Kang, H.-J., Im, S.-H., 2015. Probiotics as an immune modulator. Journal of Nutritional Science and Vitaminology 61, S103–S105.

Kang, J.G., Amar, M.J., Remaley, A.T., Kwon, J., Blackshear, P.J., Wang, P.Y., Hwang, P.M., 2011. Zinc finger protein tristetraprolin interacts with CCL3 mRNA and regulates tissue inflammation. Journal of immunology 187, 2696–2701.

Khosravi, A., Yáñez, A., Price, J.G., Chow, A., Merad, M., Goodridge, H.S., Mazmanian, S.K., 2014. Gut microbiota promote hematopoiesis to control bacterial infection. Cell host & microbe 15, 374–381.

Klaasen, H.L., Van der Heijden, P.J., Stok, W., Poelma, F.G., Koopman, J.P., Van den Brink, M.E., Bakker, M.H., Eling, W.M., Beynen, A.C., 1993. Apathogenic, intestinal, segmented, filamentous bacteria stimulate the mucosal immune system of mice. Infection and immunity 61, 303–306.

Kubinak, J.L., Petersen, C., Stephens, W.Z., Soto, R., Bake, E., O’Connell, R.M., Round, J.L., 2015. MyD88 signaling in T cells directs IgA-mediated control of the microbiota to promote health. Cell host & microbe 17, 153–163.

Kum, C., Sekkin, S., 2011. The Immune System Drugs in Fish: Immune Function, Immunoassay, Drugs.

Kumar, H., Kawai, T., Akira, S., 2011. Pathogen Recognition by the Innate Immune System. Int Rev Immunol 30, 16–34.

Kumar, S., Ingle, H., Prasad, D.V., Kumar, H., 2013. Recognition of bacterial infection by innate immune sensors. Crit Rev Microbiol 39, 229–246.

Lai, Y., Cogen, A.L., Radek, K.A., Park, H.J., Macleod, D.T., Leichtle, A., Ryan, A.F., Di Nardo, A., Gallo, R.L., 2010. Activation of TLR2 by a small molecule produced by Staphylococcus epidermidis increases antimicrobial defense against bacterial skin infections. J Invest Dermatol 130, 2211–2221.

Langfelder, P., Horvath, S., 2008. WGCNA: an R package for weighted correlation network analysis. BMC Bioinformatics 9, 559.

Langmead, B., Salzberg, S.L., 2012. Fast gapped-read alignment with Bowtie 2. Nature methods 9, 357–359.

Larsson, E., Tremaroli, V., Lee, Y.S., Koren, O., Nookaew, I., Fricker, A., Nielsen, J., Ley, R.E., Bäckhed, F., 2012. Analysis of gut microbial regulation of host gene expression along the length of the gut and regulation of gut microbial ecology through MyD88. Gut 61, 1124.

Lefebvre, T., Miambi, E., Pando, A., Diouf, M., Rouland-Lefèvre, C., 2009. Gut-specific actinobacterial community structure and diversity associated with the wood-feeding termite species, Nasutitermes corniger (Motschulsky) described by nested PCR-DGGE analysis. Insectes Sociaux 56, 269–276.

Levy, M., Kolodziejczyk, A.A., Thaiss, C.A., Elinav, E., 2017. Dysbiosis and the immune system. Nature Reviews Immunology 17, 219–232.

Li, Y., Xie, W., Li, Q., 2016. Characterisation of the bacterial community structures in the intestine of Lampetra morii. Antonie Van Leeuwenhoek 109, 979–986.

Ling, F., Steinel, N., Weber, J., Ma, L., Smith, C., Correa, D., Zhu, B., Bolnick, D., Wang, G., 2020. The gut microbiota response to helminth infection depends on host sex and genotype. The ISME journal.

Llewellyn, A., Foey, A., 2017. Probiotic modulation of innate cell pathogen sensing and signaling events. Nutrients 9.

Lohman, B.K., Weber, J.N., Bolnick, D.I., 2016. Evaluation of TagSeq, a reliable low-cost alternative for RNAseq. Molecular ecology resources 16, 1315–1321.

Love, M.I., Huber, W., Anders, S., 2014. Moderated estimation of fold change and dispersion for RNA-seq data with DESeq2. Genome biology 15, 550.

Macpherson, A.J., McCoy, K.D., 2015. Standardised animal models of host microbial mutualism. Mucosal Immunol 8, 476–486.

Manichanh, C., Rigottier-Gois, L., Bonnaud, E., Gloux, K., Pelletier, E., Frangeul, L., Nalin, R., Jarrin, C., Chardon, P., Marteau, P., Roca, J., Dore, J., 2006. Reduced diversity of faecal microbiota in Crohn’s disease revealed by a metagenomic approach. Gut 55, 205–211.

Marchesi, J.R., Adams, D.H., Fava, F., Hermes, G.D., Hirschfield, G.M., Hold, G., Quraishi, M.N., Kinross, J., Smidt, H., Tuohy, K.M., Thomas, L.V., Zoetendal, E.G., Hart, A., 2016. The gut microbiota and host health: a new clinical frontier. Gut 65, 330–339.

Maynard, C.L., Elson, C.O., Hatton, R.D., Weaver, C.T., 2012. Reciprocal interactions of the intestinal microbiota and immune system. Nature 489, 231–241.

Mazmanian, S.K., Round, J.L., Kasper, D.L., 2008. A microbial symbiosis factor prevents intestinal inflammatory disease. Nature 453, 620–625.

McMurdie, P.J., Holmes, S., 2013. phyloseq: an R package for reproducible interactive analysis and graphics of microbiome census data. PloS one 8, e61217.

Mehlitz, A., Rudel, T., 2013. Modulation of host signaling and cellular responses by Chlamydia. Cell Commun Signal 11, 90.

Milligan-Myhre, K., Small, C.M., Mittge, E.K., Agarwal, M., Currey, M., Cresko, W.A., Guillemin, K., 2016. Innate immune responses to gut microbiota differ between oceanic and freshwater threespine stickleback populations. Disease models & mechanisms 9, 187.

Nakajima, A., Vogelzang, A., Maruya, M., Miyajima, M., Murata, M., Son, A., Kuwahara, T., Tsuruyama, T., Yamada, S., Matsuura, M., Nakase, H., Peterson, D.A., Fagarasan, S., Suzuki, K., 2018. IgA regulates the composition and metabolic function of gut microbiota by promoting symbiosis between bacteria. J Exp Med 215, 2019–2034.

Navaneethan, U., Venkatesh, P.G., Shen, B., 2010. Clostridium difficile infection and inflammatory bowel disease: understanding the evolving relationship. World J Gastroenterol 16, 4892–4904.

Oh, J., Freeman, A.F., Program, N.C.S., Park, M., Sokolic, R., Candotti, F., Holland, S.M., Segre, J.A., Kong, H.H., 2013. The altered landscape of the human skin microbiome in patients with primary immunodeficiencies. Genome Res 23, 2103–2114.

Okumura, R., Takeda, K., 2018. Maintenance of intestinal homeostasis by mucosal barriers. Inflamm Regen 38, 5.

Oren, A., 2008. Microbial life at high salt concentrations: phylogenetic and metabolic diversity. Saline Systems 4, 2–2.

Org, E., Parks, B.W., Joo, J.W., Emert, B., Schwartzman, W., Kang, E.Y., Mehrabian, M., Pan, C., Knight, R., Gunsalus, R., Drake, T.A., Eskin, E., Lusis, A.J., 2015. Genetic and environmental control of host-gut microbiota interactions. Genome Res 25, 1558–1569.

Palm, N.W., de Zoete, M.R., Flavell, R.A., 2015. Immune-microbiota interactions in health and disease. Clin Immunol 159, 122–127.

Palmer-Young, E.C., Ngor, L., Burciaga Nevarez, R., Rothman, J.A., Raffel, T.R., McFrederick, Q.S., 2019. Temperature dependence of parasitic infection and gut bacterial communities in bumble bees. Environmental microbiology 21, 4706–4723.

Pedersen, H.K., Gudmundsdottir, V., Nielsen, H.B., Hyotylainen, T., Nielsen, T., Jensen, B.A., Forslund, K., Hildebrand, F., Prifti, E., Falony, G., Le Chatelier, E., Levenez, F., Dore, J., Mattila, I., Plichta, D.R., Poho, P., Hellgren, L.I., Arumugam, M., Sunagawa, S., Vieira-Silva, S., Jorgensen, T., Holm, J.B., Trost, K., Meta, H.I.T.C., Kristiansen, K., Brix, S., Raes, J., Wang, J., Hansen, T., Bork, P., Brunak, S., Oresic, M., Ehrlich, S.D., Pedersen, O., 2016. Human gut microbes impact host serum metabolome and insulin sensitivity. Nature 535, 376–381.

Qin, Y., Roberts, J.D., Grimm, S.A., Lih, F.B., Deterding, L.J., Li, R., Chrysovergis, K., Wade, P.A., 2018. An obesity-associated gut microbiome reprograms the intestinal epigenome and leads to altered colonic gene expression. Genome biology 19, 7.

Reddick, L.E., Alto, Neal M., 2014. Bacteria Fighting Back: How Pathogens Target and Subvert the Host Innate Immune System. Molecular cell 54, 321–328.

Reikvam, D.H., Erofeev, A., Sandvik, A., Grcic, V., Jahnsen, F.L., Gaustad, P., McCoy, K.D., Macpherson, A.J., Meza-Zepeda, L.A., Johansen, F.E., 2011. Depletion of murine intestinal microbiota: effects on gut mucosa and epithelial gene expression. PloS one 6, e17996.

Rennison, D.J., Rudman, S.M., Schluter, D., 2019. Parallel changes in gut microbiome composition and function during colonization, local adaptation and ecological speciation. Proceedings. Biological sciences / The Royal Society 286, 20191911.

Richards, A.L., Muehlbauer, A.L., Alazizi, A., Burns, M.B., Findley, A., Messina, F., Gould, T.J., Cascardo, C., Pique-Regi, R., Blekhman, R., Luca, F., 2019. Gut Microbiota Has a Widespread and Modifiable Effect on Host Gene Regulation. mSystems 4, e00323–00318.

Rizzatti, G., Lopetuso, L.R., Gibiino, G., Binda, C., Gasbarrini, A., 2017. Proteobacteria: A Common Factor in Human Diseases. Biomed Res Int 2017, 9351507.

Rosenberg, E., 2014. The Family Chitinophagaceae, in: Rosenberg, E., DeLong, E.F., Lory, S., Stackebrandt, E., Thompson, F. (Eds.), The Prokaryotes: Other Major Lineages of Bacteria and The Archaea. Springer Berlin Heidelberg, Berlin, Heidelberg, pp. 493–495.

Rosser, E.C., Oleinika, K., Tonon, S., Doyle, R., Bosma, A., Carter, N.A., Harris, K.A., Jones, S.A., Klein, N., Mauri, C., 2014. Regulatory B cells are induced by gut microbiota-driven interleukin-1beta and interleukin-6 production. Nat Med 20, 1334–1339.

Rosshart, S.P., Vassallo, B.G., Angeletti, D., Hutchinson, D.S., Morgan, A.P., Takeda, K., Hickman, H.D., McCulloch, J.A., Badger, J.H., Ajami, N.J., Trinchieri, G., Pardo-Manuel de Villena, F., Yewdell, J.W., Rehermann, B., 2017. Wild Mouse Gut Microbiota Promotes Host Fitness and Improves Disease Resistance. Cell 171, 1015–1028.e1013.

Rowland, I., Gibson, G., Heinken, A., Scott, K., Swann, J., Thiele, I., Tuohy, K., 2018. Gut microbiota functions: metabolism of nutrients and other food components. Eur J Nutr 57, 1–24.

Rusconi, B., Greub, G., 2011. Chlamydiales and the innate immune response: friend or foe? FEMS Immunol Med Microbiol 61, 231–244.

Russell, J.T., Roesch, L.F.W., Ordberg, M., Ilonen, J., Atkinson, M.A., Schatz, D.A., Triplett, E.W., Ludvigsson, J., 2019. Genetic risk for autoimmunity is associated with distinct changes in the human gut microbiome. Nature communications 10, 3621.

Salzman, N.H., 2010. Paneth cell defensins and the regulation of the microbiome: detente at mucosal surfaces. Gut microbes 1, 401–406.

Scher, J.U., Abramson, S.B., 2011. The microbiome and rheumatoid arthritis. Nat Rev Rheumatol 7, 569–578.

Schloss, P.D., Westcott, S.L., Ryabin, T., Hall, J.R., Hartmann, M., Hollister, E.B., Lesniewski, R.A., Oakley, B.B., Parks, D.H., Robinson, C.J., Sahl, J.W., Stres, B., Thallinger, G.G., Van Horn, D.J., Weber, C.F., 2009. Introducing mothur: open-source, platform-independent, community-supported software for describing and comparing microbial communities. Applied and environmental microbiology 75, 7537–7541.

Schroeder, B.O., 2019. Fight them or feed them: how the intestinal mucus layer manages the gut microbiota. Gastroenterol Rep (Oxf) 7, 3–12.

Shokryazdan, P., Faseleh Jahromi, M., Navidshad, B., Liang, J.B., 2017. Effects of prebiotics on immune system and cytokine expression. Med Microbiol Immunol 206, 1–9.

Slobodkin, A., 2014. The Family Peptostreptococcaceae, in: Rosenberg, E., DeLong, E.F., Lory, S., Stackebrandt, E., Thompson, F. (Eds.), The Prokaryotes: Firmicutes and Tenericutes. Springer Berlin Heidelberg, Berlin, Heidelberg, pp. 291–302.

Smith, C.C., Snowberg, L.K., Gregory Caporaso, J., Knight, R., Bolnick, D.I., 2015. Dietary input of microbes and host genetic variation shape among-population differences in stickleback gut microbiota. The ISME journal 9, 2515–2526.

Smith, K., McCoy, K.D., Macpherson, A.J., 2007. Use of axenic animals in studying the adaptation of mammals to their commensal intestinal microbiota. Seminars in immunology 19, 59–69.

Sommer, F., Nookaew, I., Sommer, N., Fogelstrand, P., Backhed, F., 2015. Site-specific programming of the host epithelial transcriptome by the gut microbiota. Genome biology 16, 62.

Tanekhy, M., Matsuda, S., Itano, T., Kawakami, H., Kono, T., Sakai, M., 2010. Expression of cytokine genes in head kidney and spleen cells of Japanese flounder (Paralichthys olivaceus) infected with Nocardia seriolae. Vet Immunol Immunopathol 134, 178–183.

Tanoue, T., Morita, S., Plichta, D.R., Skelly, A.N., Suda, W., Sugiura, Y., Narushima, S., Vlamakis, H., Motoo, I., Sugita, K., Shiota, A., Takeshita, K., Yasuma-Mitobe, K., Riethmacher, D., Kaisho, T., Norman, J.M., Mucida, D., Suematsu, M., Yaguchi, T., Bucci, V., Inoue, T., Kawakami, Y., Olle, B., Roberts, B., Hattori, M., Xavier, R.J., Atarashi, K., Honda, K., 2019. A defined commensal consortium elicits CD8 T cells and anti-cancer immunity. Nature 565, 600–605.

Thaiss, C.A., Zmora, N., Levy, M., Elinav, E., 2016. The microbiome and innate immunity. Nature 535, 65–74.

Vaziri, N.D., Wong, J., Pahl, M., Piceno, Y.M., Yuan, J., DeSantis, T.Z., Ni, Z., Nguyen, T.H., Andersen, G.L., 2013. Chronic kidney disease alters intestinal microbial flora. Kidney Int 83, 308–315.

Walke, J.B., Becker, M.H., Loftus, S.C., House, L.L., Teotonio, T.L., Minbiole, K.P., Belden, L.K., 2015. Community Structure and Function of Amphibian Skin Microbes: An Experiment with Bullfrogs Exposed to a Chytrid Fungus. PloS one 10, e0139848.

Wang, S., Ye, Q., Zeng, X., Qiao, S., 2019. Functions of Macrophages in the Maintenance of Intestinal Homeostasis. J Immunol Res 2019, 1512969.

Wanke, I., Steffen, H., Christ, C., Krismer, B., Gotz, F., Peschel, A., Schaller, M., Schittek, B., 2011. Skin commensals amplify the innate immune response to pathogens by activation of distinct signaling pathways. J Invest Dermatol 131, 382–390.

Weber, J.N., Kalbe, M., Shim, K.C., Erin, N.I., Steinel, N.C., Ma, L., Bolnick, D.I., 2017a. Resist globally, infect locally: A transcontinental test of adaptation by stickleback and their tapeworm parasite. American Naturalist 189, 43–57.

Weber, J.N., Steinel, N.C., Shim, K.C., Bolnick, D.I., 2017b. Recent evolution of extreme cestode growth suppression by a vertebrate host. Proc Natl Acad Sci U S A 114, 6575–6580.

Winans, N.J., Walter, A., Chouaia, B., Chaston, J.M., Douglas, A.E., Newell, P.D., 2017. A genomic investigation of ecological differentiation between free-living and Drosophila-associated bacteria. Molecular ecology 26, 4536–4550.

Wright, R.M., Aglyamova, G.V., Meyer, E., Matz, M.V., 2015. Gene expression associated with white syndromes in a reef building coral, Acropora hyacinthus. BMC genomics 16, 371.

Xu, Z., Takizawa, F., Casadei, E., Shibasaki, Y., Ding, Y., Sauters, T.J.C., Yu, Y., Salinas, I., Sunyer, J.O., 2020. Specialization of mucosal immunoglobulins in pathogen control and microbiota homeostasis occurred early in vertebrate evolution. Sci Immunol 5.

Zhang, Z., Tang, H., Chen, P., Xie, H., Tao, Y., 2019. Demystifying the manipulation of host immunity, metabolism, and extraintestinal tumors by the gut microbiome. Signal Transduct Target Ther 4, 41.

Zhao, Q., Elson, C.O., 2018. Adaptive immune education by gut microbiota antigens. Immunology 154, 28–37.

Zheng, D., Liwinski, T., Elinav, E., 2020. Interaction between microbiota and immunity in health and disease. Cell Res 30, 492–506.

Zvanych, R., Lukenda, N., Kim, J.J., Li, X., Petrof, E.O., Khan, W.I., Magarvey, N.A., 2014. Small molecule immunomodulins from cultures of the human microbiome member Lactobacillus plantarum. J Antibiot (Tokyo) 67, 85–88.

